# Implicit perceptual memory can increase or decrease with ageing

**DOI:** 10.1101/736579

**Authors:** KA Zhivago, Sneha Shashidhara, Ranjini Garani, Simran Purokayastha, Naren P. Rao, Aditya Murthy, SP Arun

**Affiliations:** Centre for Neuroscience, Indian Institute of Science, Bangalore, India; Department of Psychiatry, National Institute of Mental Health and Neurosciences, Bangalore, India

## Abstract

A decline in declarative or explicit memory has been extensively characterized in cognitive ageing and is a hallmark of cognitive impairments. However, whether and how implicit perceptual memory varies with ageing or cognitive impairment is unclear. Here, we compared implicit perceptual memory and explicit memory measures in three groups of subjects: (1) 59 healthy young volunteers (20-30 years); (2) 238 healthy old volunteers (50-90 years) and (3) 21 patients with mild cognitive impairment MCI (50-90 years). To measure explicit memory, subjects were tested on standard recognition and recall tasks. To measure implicit perceptual memory, we used a classic perceptual priming paradigm. Subjects had to report the shape of a visual search pop-out target. Implicit priming was measured as the speedup in response time for targets with the same vs different color/position on consecutive trials.

Our main findings are as follows: (1) Explicit memory was weaker in old compared to young subjects, and in MCI compared to age-matched controls; (2) Surprisingly, implicit perceptual memory did not always decline with age: color priming was smaller in older subjects but position priming was larger; (3) Position priming was less frequent in the MCI group compared to age-matched controls; (4) Implicit and explicit memory measures were uncorrelated in all three groups. Thus, implicit memory can increase or decrease with age or cognitive impairment, but this decline does not covary with explicit memory. We propose that incorporating explicit and implicit measures can yield a richer characterization of memory.

## INTRODUCTION

Memory has broadly been classified into explicit and implicit memory (Squire, 1992; Gazzaniga, Ivry and Mangun, 2014). Explicit memory is consciously accessible and declarative; it is measured by how well subjects can recall items that were previously studied (Strauss, Sherman and Spreen, 2006). By contrast, implicit memory is unconscious and non-declarative; it is measured by the facilitation in the response to previously experienced items (Fleischman *et al.*, 2005; Spataro *et al.*, 2016). Explicit and implicit memory are widely thought to be dissociable (Gazzaniga, Ivry and Mangun, 2014). Evidence in favour of this account comes from the fact that explicit memory is impaired in amnesic patients but not implicit memory (Graf, Squire and Mandler, 1984; Fleischman *et al.*, 2005; Fleischman, 2007; Gazzaniga, Ivry and Mangun, 2014), although the opposite outcome – impaired implicit and intact explicit memory – has been observed only rarely (Gabrieli *et al.*, 1995; Ward, Berry and Shanks, 2013; Gazzaniga, Ivry and Mangun, 2014). This leaves open the possibility that both explicit and implicit memory probe a common memory system in qualitatively different ways (Ward, Berry and Shanks, 2013).

Since a decline in explicit memory is a hallmark of both ageing and cognitive disorders, the natural question is whether implicit memory is also affected (Ward, Berry and Shanks, 2013). Previous studies have addressed this question by comparing explicit and implicit memory measures across age and in cognitive impairments (Fleischman, 2007). Implicit memory in these studies was typically measured as an increased probability of producing a studied item in an unrelated task, or the facilitated recognition of a fragmented picture after it was previously viewed. The results are mixed: explicit memory clearly declines with age and between patients and controls, but implicit memory declines in some cases (Perri *et al.*, 2007; Ballesteros, Mayas and Reales, 2013; Boccia, Silveri and Guariglia, 2014) but not others (Fleischman *et al.*, 2005). It has been proposed that an implicit memory deficit could be an early sign for the onset of dementia (Fleischman, 2007). However, these implicit memory tests have several problems. First, subjects may use explicit memory during the study phase. This “explicit contamination” can be mitigated but is extremely tricky to fully rule out (Fleischman, 2007). Second, these tests assume a minimum proficiency in verbal and object naming which may vary widely especially in diverse populations with varying degrees of multilingualism and literacy. One potential solution to these issues is to develop tasks that are culture-free with no study phase.

Here, we devised an implicit memory paradigm based on a classic perceptual priming effect, known as priming of pop-out (Maljkovic and Nakayama, 1994, 1996). In this paradigm, subjects are faster to respond to the shape of a pop-out target when its color or position is repeated. This task has two key advantages over previously used implicit memory measures. First, the task is easy to comprehend and assumes no prior knowledge of objects or words, making it suitable for use on diverse populations without placing inordinate demands on verbal knowledge. Second, the task does not involve a study phase thereby mitigating any explicit contamination.

## METHODS

All subjects had normal or corrected-to-normal vision and gave written informed consent to an experimental protocol approved by the Institutional Human Ethics Committee of the Indian Institute of Science, Bangalore. All subjects were compensated monetarily for their participation.

### Subjects

Young volunteers were all students from the campus. The older volunteers and patients were literate and from an urban environment, and were participants of an ongoing Tata Longitudinal Study of Aging. In all, we analysed data from 59 young subjects (25 ± 3 years, 29 female; years of education: 19 ± 3 years), 238 old subjects (66 ± 8 years, 108 female; years of education: 17 ± 3 years), and 21 MCI patients (72 ± 10 years, 4 female; years of education: 15 ± 5 years). From these older subjects, we selected a subset of participants whose age matched with one of the MCI patients. This age-matched control group comprised 92 subjects (69 ± 8 years, 39 female; years of education: 17 ± 3 years). The older volunteers were large in number because they were participants of an ongoing longitudinal study of healthy volunteers. By contrast, the younger volunteers group were recruited exclusively for the purpose of the cross-sectional comparisons in the present study.

### MCI Diagnosis

A trained neurologist or psychiatrist administered the Clinical Dementia rating (CDR) scale (Hughes et al., 1982) and a diagnosis was given based upon the scores. CDR is administered by interviewing the volunteer and their primary caregiver (typically a close relative), which took about 20-30 mins. A score of 0 corresponded to a healthy volunteer and 0.5 to MCI.

### Global cognitive scores

To validate the MCI Diagnosis obtained from the CDR scale, we also compared the global cognitive score ACE III [1], measured from the older volunteers as part of the Tata Longitudinal Study of Aging. As expected, ACE-III scores were significantly higher for age-matched compared to MCI (average ACE-III score: 94±4 for age-matched, 87±7 for patients, p < 0.00005, rank-sum test).

### Stimuli

All tasks were administered through HTML5/Javascript scripts run on an internet browser on a desktop computer (24-inch, width × height: 53.3 × 30.0 cm, 1920 × 1080 pixels) or laptop computer (15.4-inch, width × height: 33.0 × 20.6 cm, 2880 × 1800 pixels). All displayed items were scaled using the monitor size and viewing distance so that they subtended the same visual angle on the eye. The browser-based setup was validated by comparing it with visual search tasks written in Matlab and Psychtoolbox.

### Procedure

Subjects were seated with a computer monitor placed at a distance of ∼50 cm. Each subject performed a perceptual priming task, an object recognition task, a word recall and word recognition task in that order, as detailed below.

### Perceptual priming task

Each trial started with a gray fixation square (0.4°x0.4°) displayed for 750 ms at the center of the screen, followed by a hexagonal search array (with items placed 6° from the center) containing one oddball item among five other distractors (Figure 1A). Each target or distractor shape was a diamond measuring 2° along the longer dimension with a 1° vertical cut on the left or right side. The array comprised a red target among green distractors or vice-versa, with the target chosen to appear randomly either at the leftmost or the rightmost location. Subjects were instructed to indicate whether the oddball target diamond was cut on the right or left side by pressing the corresponding arrow key on a keyboard. The search array was displayed for 10 seconds, or until the subject made a valid response, whichever was shorter. Target color and position were counterbalanced (i.e. equal numbers of red/green × left/right trials) and presented in random order. Error trials and trials with no response were repeated later after a random number of other trials. Subjects performed a total of 200 correct trials.

**Figure 1.**
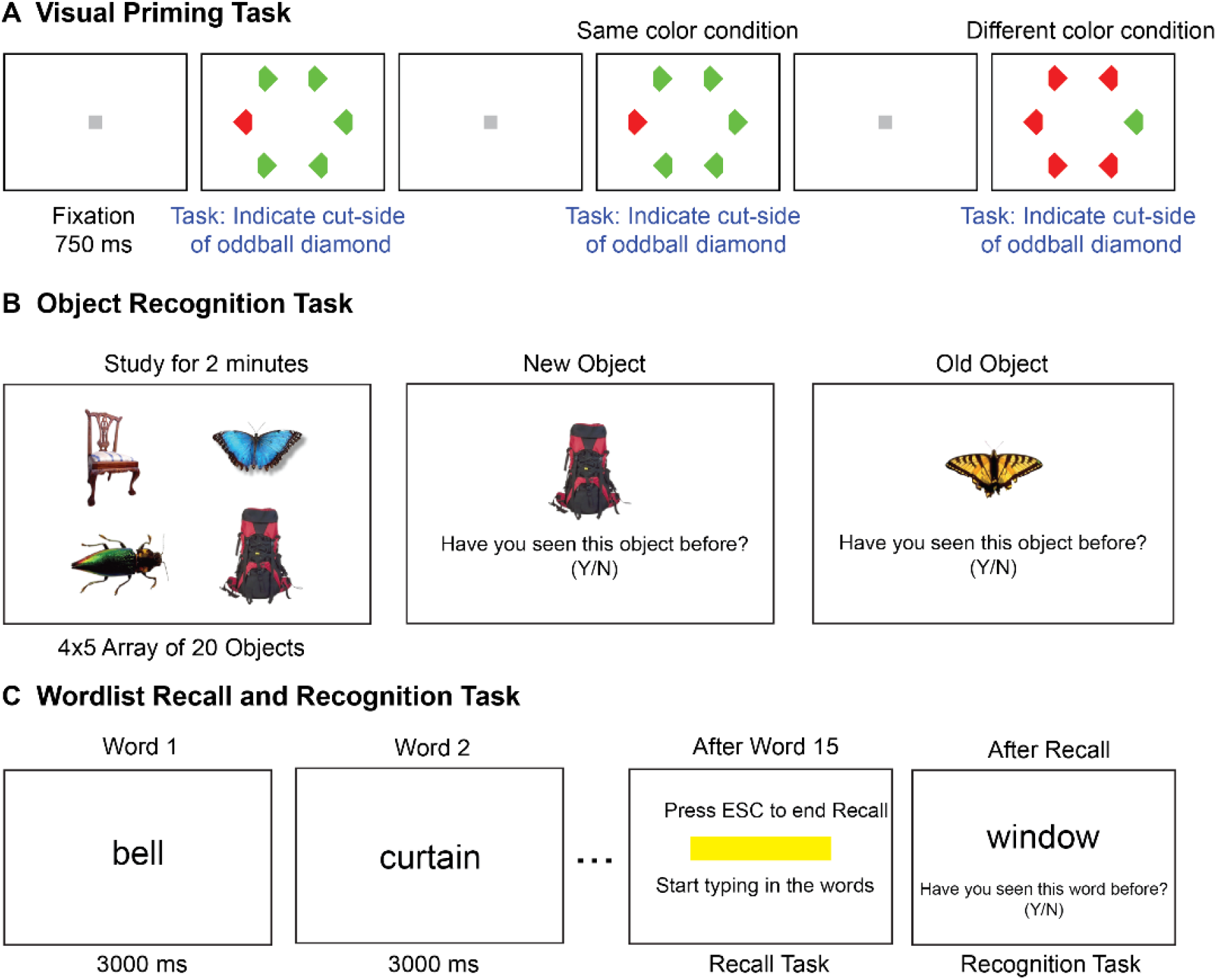
Implicit and explicit memory tasks. (A) Schematic of the visual priming task with its two possible conditions. Each trial started with a gray fixation square followed by a search array. The second trial shows the odd–colored target with the same color (*red*) as the previous target, hence it is the same-color condition. The next trial shows the case where the target color (*green*) is different from the previous trial (*red*), hence it is the different-color condition. Subjects make faster responses on same-color trials compared to different-color trials, indicative of an implicit memory. (B)Schematic of the object recognition task. Subjects were asked to study a set of 20 pictures for 2 minutes. In the test phase (second & third panels), one picture was shown at a time and subjects had to indicate whether the object was shown or not shown during the study phase. (C) Schematic of the word recall and recognition tasks. During the study phase (*left panel*), 15 words are presented for 3 seconds each in a sequence. In the recall phase (*middle panel*), a text box appeared on the screen, and the experimenter typed in the words recalled verbally by the subject. In the recognition task (*right panel*), 30 words were presented in sequence, and subjects had to indicate whether the word was shown or not during the study phase.

Priming strength was calculated as the percent decrease in response time on a trial preceded by a target of the same color/position relative to the response time on trials where the target was of a different color/position. Thus,

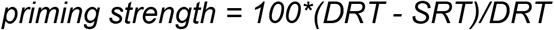

where*, DRT* is the mean reaction time of trials preceded by a different color/position compared to the current trial and *SRT* is the mean reaction time of trials preceded by the same color/position compared to the current trial. In the analyses reported here, we calculated priming strength only using consecutive trials with correct responses, but we obtained qualitatively similar results upon using all trials regardless of error status. We also obtained qualitatively similar results on using SRT in the denominator instead of DRT.

### Object recognition task

Each subject was shown a total of 20 objects (measuring 3.5° along the longer dimension) arranged in a 4 × 5 grid for 2 minutes. These were pictures of animals, household objects, vehicles, etc. selected from Hemera Photo Objects (copyright 2015, corresponding author) (Figure 1B). After the study phase, 40 pictures were presented, one at a time, prompting the subject to press ‘y’ or ‘n’ key to indicate whether (s)he saw the object during the study phase or not, with no time restriction for each response. A total of 20 of the 40 pictures were from the memorization phase and the remaining 20 new pictures were drawn from the same basic-level category as the 20 old pictures. The subjects were instructed that they do not have to name any picture or for that matter give any descriptive account. It was designed to be a purely visual test. Since our cohort was urban and literate, all the pictures were culture-fair and we have no reason to suspect novelty to affect the recognition performance in any way.

Object recognition memory performance was characterized for each subject by calculating the total percentage correct across old and new items. We observed no consistent differences in response bias (towards old or new) across the four groups of subjects.

### Wordlist recall and recognition task

Each subject performed a word recall block and a recognition block, always in this order (Figure 1C). Each recall block started with a study phase in which 15 words (e.g. colour, garden, coffee, house, etc.) were presented on the screen in a predefined sequence, with each word shown for 3 seconds. This was followed by a recall phase in which the subjects were asked to recall as many words as they could from the study phase, and the experimenter typed in the words. Subjects were free to recall words in any order and take any amount of time. The block was stopped once the subject declared that they could not remember any more words. Word recall performance was calculated for each subject as the fraction of words correctly recalled out of the full list.

During the recognition block, one word was presented at a time and the subject was asked to report with a key press, if the word was presented in the recall block (‘y’ for yes and ‘n’ for no). A total of 30 words were presented, 15 of which belonged to the recall block and 15 were new. The new words were drawn from similar categories as the study words (e.g. crayon, tree, home, etc.). Word recognition performance was calculated for each subject as the fraction of old and new words that elicited a correct response. We observed no consistent difference in response bias towards old or new words across subject groups. Because our cohort was urban and literate, we did not expect word novelty affecting recall or recognition.

The word tasks above involved making computer responses, in contrast to the standard word tasks in which there are multiple study phases and the subject is asked to make only verbal responses. To assess this possibility, we compared our word task scores with the verbally administered word recall and recognition tasks from the <anonymized> Longitudinal Study, which were always administered before our word tasks but using an independent word list. This revealed a significant positive correlation (immediate word recall: r = 0.37, p < 0.000005 and immediate word recognition: r = 0.35, p < 0.000005 across all older volunteers). We also did not find any systematic relation between the number of years of education and the explicit memory measures.

### Statistical group comparisons

We used non-parametric tests to compare statistics between subject groups (Wilcoxon signed-rank test for paired comparisons and Wilcoxon rank-sum test for unpaired comparisons), and ANOVA in multifactorial comparisons. Using parametric or nonparametric tests in general yielded qualitatively similar levels of significance.

## RESULTS

Our goal was to characterize explicit and implicit memory across age and cognitive impairments. We compared these measures between young vs older volunteers and between MCI patients and age-matched controls. The tasks performed by each subject are detailed in Figure 1.

### Perceptual priming task

All four groups of subjects were highly accurate on this task (accuracy mean ± sd: 96±5% for young, 97±4% for old; 97±4% for age-matched controls, and 95±7% for patients). Compared to young subjects, older subjects were generally more accurate (p < 0.0005, rank-sum test on overall accuracy for young vs old), but were considerably slower (average RT, mean±sd: 1.02±0.34 s for young; 1.75±0.56 s for old; p < 0.00005, rank-sum test). Likewise compared to age-matched controls, patients were equally accurate (p = 0.06, rank-sum test) and fast (average RT: 1.97±0.62 s, 1.84±0.62 s, p = 0.33, rank-sum test).

To measure implicit memory, we compared the reaction time on trials preceded by a target of the same versus different color or position. In all four groups (young, old, age-matched controls and patients), subjects were faster for same-color trials compared to different-color trials in all groups (Figure 2A), and were faster for same-position trials compared to different-position trials (Figure 2D). A detailed statistical comparison is provided in Section S1. Thus, priming of pop-out is robustly present at the group level in young, old, age-matched controls and patients.

**Figure 2:**
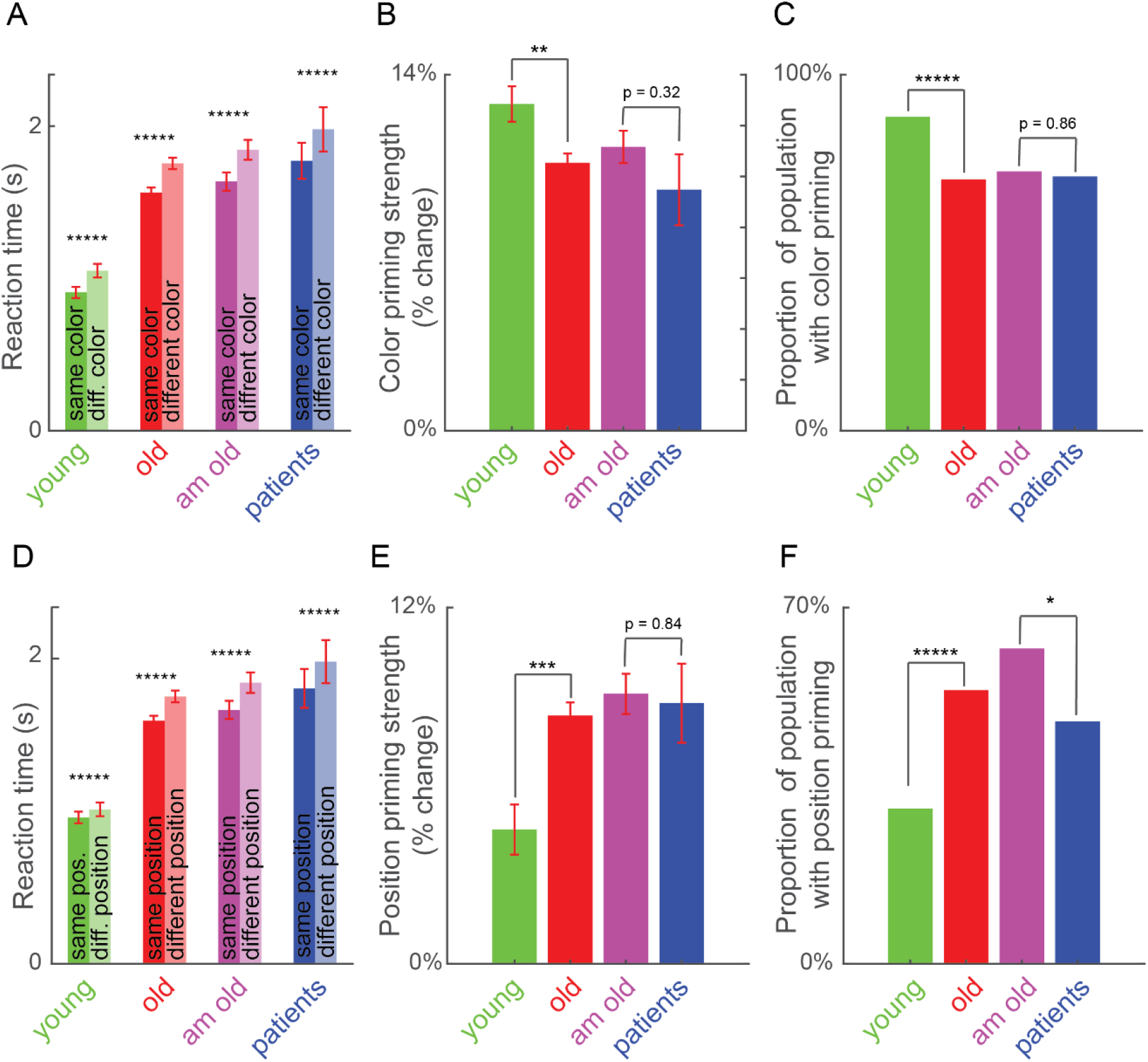
Perceptual priming effects across age and cognitive impairments. (A) Average reaction time for same-color (dark) and different-color (light) trials across young, old, age-matched controls (am old) and patients. Asterisks indicate statistical significance of the main effect of color in an ANOVA (Section S2): * is p < 0.05, ** is p < 0.005 etc. Error bars indicate s.e.m across subjects. (B) Color priming strength across groups. Asterisks indicate statistical significance comparing priming strength across subjects between groups, using a rank-sum test (* is p < 0.05, ** is p < 0.005 etc). Error bars indicate s.e.m across subjects. (C) Proportion of population with significant color priming shown for each group of subjects. Asterisks indicate statistical significance comparing the rate of incidence between each pair of groups, using a chi-squared test, with conventions as before. (D-F) Same as A-C but for position priming.

### Do color and position priming vary with age or with cognitive impairment?

Next we asked whether the strength of priming was different across age or with cognitive impairment. To this end, we calculated the priming strength for each subject as the percentage change in reaction time between primed (i.e. same color or position) and unprimed (different color or position) trials. Calculating the percentage change ensures that the measure is normalized to the speed of each subject and therefore comparable across subject groups. To ascertain the reliability of priming strength we calculated priming strength using the first and second half of the trials for each subject, and asked whether the two measures were correlated across subjects. This split-half correlation, which is an index of reliability, was strikingly high for all groups (r = 0.90, 0.89, 0.92 and 0.92 for young, old, patients and age-matched, p < 0.000005 for all groups).

Color priming strength was significantly weaker for older subjects compared to young subjects (Figure 2B; Priming strength: 13±1% for young, 11±0% for old, p < 0.05, rank-sum test). It was weaker for patients compared to age-matched controls, but this effect was not statistically significant (Figure 2B; Priming strength: 9±1% for patients, 11±1% for age-matched controls, p = 0.32, rank-sum test). By contrast, position priming was stronger in older subjects compared to young (Figure 2E; Priming strength: 5±1% for young, 8±0% for old, p < 0.0005, rank-sum test). As with color priming, position priming was numerically weaker in patients compared to age-matched controls but this effect was not statistically significant (Priming strength: 9.1±0.7% for patients, 8.8±1.3% for age-matched controls, p = 0.84, rank-sum test).

The variations in color and priming strength between young and old subjects may indicate a general change in priming strength with age, or a selective change in a subset of subjects. To explore these possibilities, we performed a subject-wise analysis to detect the presence of color or position priming. We performed an ANOVA on the response times of each subject, with color, position and motor priming as factors. The fraction of subjects with a significant color priming effect in each group is shown in Figure 2C. Color priming was less prevalent in old compared to young subjects (Figure 2C; Percentage of subjects with significant color priming: 88% for young, 71% for old, p < 0.000005, chi-squared test comparing young subjects with and without color priming against the numbers expected from the incidence seen in the larger group i.e. old subjects). It was also less prevalent among patients compared to age-matched controls, but this difference was not statistically significant (Figure 2C; Percentage of subjects with significant color priming: 71% for patients, 73% for age-matched controls, p = 0.86, chi-squared test).

By contrast, position priming was more prevalent among old compared to young subjects (Figure 2F; Percent of subjects with significant position priming: 31% for young, 54% for old, p < 0.000005, chi-squared test). It was also significantly less prevalent among MCI compared to age-matched controls (Figure 2F; Percentage of subjects with significant position priming: 48% for patients, 62% for age-matched controls, p < 0.05, chi-squared test).

In sum, we conclude that implicit perceptual priming for color and position show differential effects across age: Older subjects show weaker color priming but stronger position priming. Position priming was less prevalent among MCI patients compared to age-matched controls. By contrast motor priming, as measured by the speedup in making the same vs different motor response across consecutive trials, showed only weak differences (Section S1).

### Explicit memory differences

We observed robust differences in explicit memory between the two groups. Object recognition memory was significantly weaker in old compared to young subjects (Figure 3A; average accuracy: 92±1% for old, 96±1 for young, p < 0.05, rank-sum test) and in patients compared to age-matched controls (Figure 2A; average accuracy: 84±1% for patients, 91±1 for age-matched controls, p < 0.005, rank-sum test). Likewise, word recall was significantly worse for old compared to young subjects (Figure 2B; average percentage of words recalled: 45±1% for old, 55±2 for young, p < 0.0005, rank-sum test). Patients showed weaker word recall compared to age-matched controls, but this effect was not significant (average percentage of words recalled: 39±4% for patients, 44±2 for age-matched controls, p = 0.17, rank-sum test). Finally, word recognition was weaker for old compared to young subjects (Figure 2C; average accuracy on old/new word recognition: 87±1% for old, 92±2 for young, p < 0.005, rank-sum test). Patients showed significantly worse word recognition to controls (Figure 2C; average accuracy: 80±2% for patients, 87±1 for age-matched controls, p < 0.05, rank-sum test). We obtained qualitatively similar trends using other measures (d’ or hits minus false alarms).

**Figure 3:**
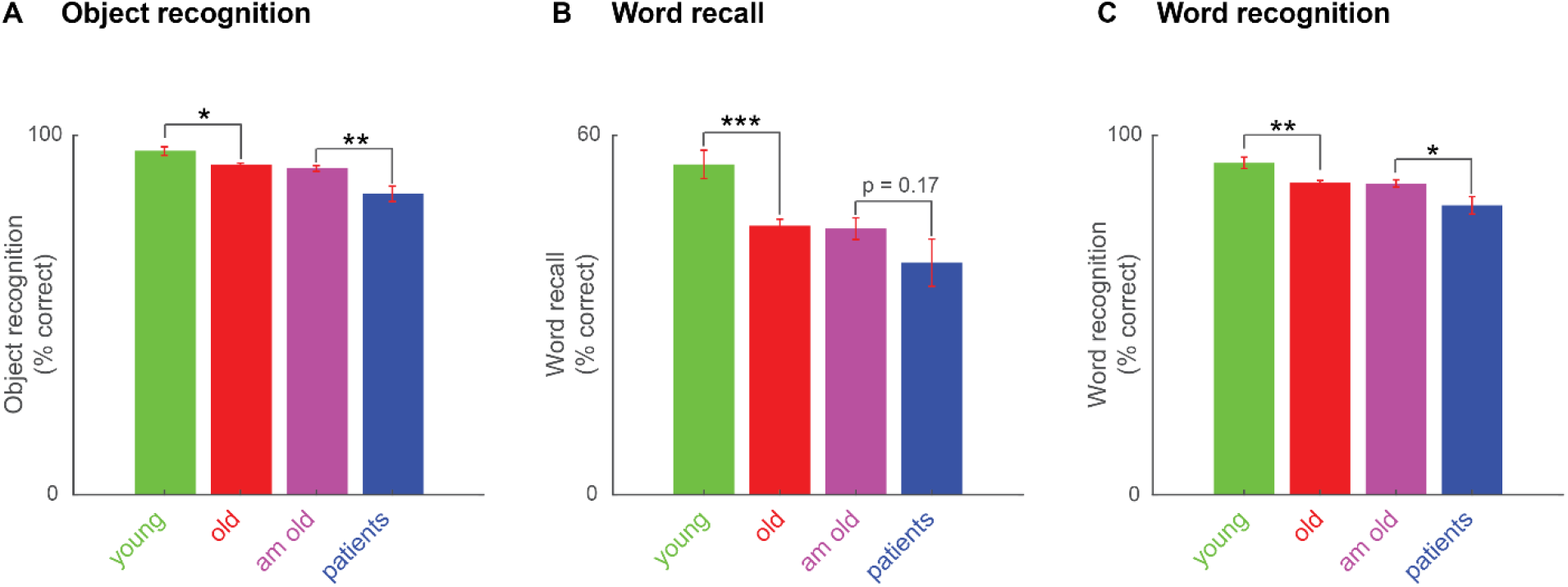
Explicit memory variations across age and cognitive impairments. (A) Object recognition accuracy (percentage correct) across all subject groups. Asterisks indicate statistical significance comparing subject-wise accuracy using a rank-sum test. Conventions are as before. (B) Word recall performance (percentage correct) across all four subject groups, with conventions as in A. (C) Word recognition performance (percentage correct) across all four subject groups, with conventions as in A.

### Relation between implicit and explicit memory

To investigate whether explicit and implicit memory covaried across subjects, we calculated the pairwise correlation between implicit and explicit measures for each group. The resulting correlations are shown in Figure 4. We observed significant correlations only in older subjects, presumably because this was the group with the largest numbers. In the older subject group, explicit memory measures were all highly correlated (Figure 4B). Color and position priming strength were not significantly correlated (r = 0.08, p = 0.22). Importantly there was no significant correlation between explicit and implicit memory measures (Figure 4B). We conclude that implicit and explicit memory are unrelated in their variation across subjects.

**Figure 4.**
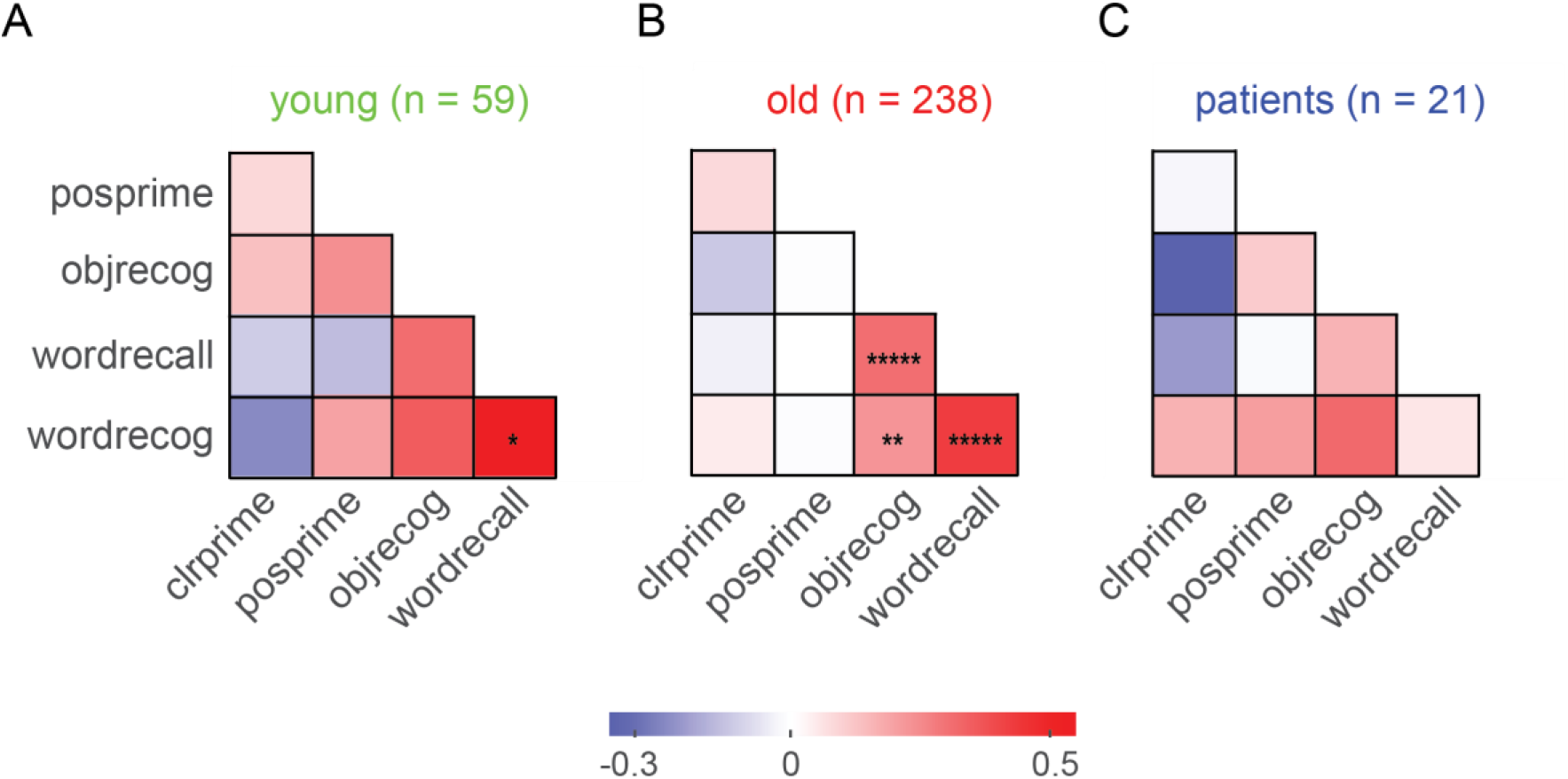
Covariation in explicit and implicit memory across subjects. (A) Pairwise correlation between explicit and implicit memory measures for young subjects. Legends: *clrprime*: color priming strength, *posprime*: position priming strength, *objrecog*: object recognition accuracy, *wordrecall*: word recall accuracy, *wordrecog*: word recognition accuracy. Each entry indicates the Pearson’s correlation coefficient between a given pair of memory measures across subjects. Asterisks indicate statistical significance of these correlations (*is p < 0.05, ** is 0.005 etc). (B) Same as A, but for older subjects. (C) Same as A, but for MCI patients.

## DISCUSSION

Here we characterized implicit memory using perceptual priming and explicit memory across age and cognitive impairments. Our main findings are: (1) Color priming is weaker but position priming is stronger in older subjects, suggesting that perceptual priming can increase or decrease with age; (2) Perceptual priming and explicit memory are uncorrelated across subjects; (3) MCI patients show weaker perceptual position priming compared to age-matched controls. Below we discuss these findings in the context of the existing literature.

Our finding that color priming is weaker but position priming is stronger in older subjects suggests that these two kinds of perceptual priming vary differently with age. Both observations are novel to the best of our knowledge, since the vast majority of implicit memory studies have focused on implicit verbal priming of previously studied items (Fleischman, 2007). The finding that color priming is weaker in older subjects is consistent with a decline in repetition priming observed previously (Gordon *et al.*, 2013; Caballero, Reales and Ballesteros, 2018). However, our finding that position priming is stronger in older subjects is noteworthy since most cognitive measures decline with age. We also note that position priming was less prevalent among MCI subjects, suggesting that the presence or absence of priming can be a useful indicator of pathological aging.

The differences between color and position priming suggest that these two kinds of priming have different underlying mechanisms. Orienting to a repeated color requires spatial attention whereas orienting to the same position requires overcoming inhibition of return (Posner, 2016). Inhibition of return is a process or mechanism by which processing of recently detected stimuli are suppressed, which thereby may make attentional resources available for processing novel stimuli. The increase in position priming in older subjects can then be explained by a loss of inhibitory control with age (Lawrence, Edwards and Goodhew, 2018). This results in decreased inhibition of return, making it is easier to orient to a target repeated at the same position. By contrast, orienting to a repeated color target is likely unaffected by inhibitory mechanisms, but may be affected by a decline in attention with age or by cognitive impairments.

In sum, our results suggest that perceptual priming measures could serve as useful markers for memory impairments. We propose that both explicit and implicit memory should be characterized in order to obtain a richer characterization of memory in cognitive testing.

## Supporting information

Supplementary Analyses

## ACKNOWLEDGEMENTS

We are grateful to Profs. Vijayalakshmi Ravindranath & David Bennett for valuable suggestions. We thank Aakash Agrawal, Saurabh Farkya and other members of the Vision Lab IISc for assistance with collecting data. This study was part of the Tata Longitudinal Study of Aging funded by the Tata Trusts, with NR, AM & SPA as co-PIs and KAZ, SS, RGR and SP as research staff.

