## Supplementary Analyses for "Implicit perceptual memory can increase or decrease with ageing"

### CONTENTS


**Section S1. Supplementary Analyses**

### Section S1. Supplementary Analyses

#### Statistical analyses for implicit priming effects within each group

To assess the statistical significance of the perceptual priming effects at the group level, we performed an ANOVA on the response times with subject (number of levels equal to the number of subjects in the given group), color priming (2 levels: same/different target color compared to previous trial), position priming (2 levels: same/different target position from previous trial) and motor priming (2 levels: same/different response key pressed compared to previous trial) as factors. A main effect of subject would indicate individual differences in mean response time across subjects. A main effect of color or position priming were of interest to us because they indicate the presence of implicit memory. A main effect of motor priming would indicate faster responses when the same response has to be made on successive trials.

In the young population, we observed significant main effects of subject ($F = 190.2$, $df = 58$, $p < 0.000005$), color ($F = 595.3$, $df = 1$, $p < 0.000005$), position ($F = 87.7$, $df = 1$, $p < 0.000005$), and also significant interaction effects between subject-color, subject-color-position and color-position ($p < 0.000005$ in all cases; all other interactions were $p > 0.05$). Among older subjects, we observed significant main effects of subject ($F = 206.1$, $df = 237$, $p < 0.000005$), color ($F = 1555.7$, $df = 1$, $p < 0.000005$), and position ($F = 1076.7$, $df = 1$, $p < 0.000005$), as well as subject-color, subject-position, subject-color-position, color-position, and position-motor interaction effects ($p < 0.000005$ in these cases; all other interactions were $p > 0.05$). In the age-matched population, we observed main effects of subject ($F = 233.8$, $df = 91$, $p < 0.000005$), color ($F = 676.2$, $df = 1$, $p < 0.000005$), and position ($F = 480.9$, $df = 1$, $p < 0.000005$), as well as subject-color, subject-position, subject-color-position and color-position interactions ($p < 0.000005$ in these cases; all other effects were $p > 0.05$). Among MCI patients, we observed significant main effects of subject ($F = 195.4$, $df = 20$, $p < 0.000005$), color ($F = 138.5$, $df = 1$, $p < 0.000005$), and position ($F = 89.9$, $df = 1$, $p < 0.000005$), as well as subject-color, subject-position, subject-color-position and color-position ($p < 0.005$ in these cases; all other effects were $p > 0.05$). A post-hoc analysis of the color-position interactions revealed that the interactions did not reverse the direction of the effect but only served to enhance it. A post-hoc analysis of the subject-related interactions revealed that this was due to variability in the incidence of these effects across subjects.

#### Do subjects with significant priming also differ across groups?

Given that the prevalence of perceptual priming can change with age, we wondered whether the subjects with significant priming effects in each group also showed a difference in priming strength across groups. To investigate this issue, we compared color and position priming strength between old and young subjects that showed significant priming effects. Color priming was slightly weaker in old compared to young subjects but this effect was not significant (color priming strength: 13±0% for 168 old subjects, 14±1% for 52 young subjects, $p = 0.35$ rank-sum test). Likewise, it did not differ between patients and age-matched controls with significant effects (color priming strength: 12±1% for 15 patients, 14±1% for 67 age-matched controls, $p = 0.20$, rank-sum test). Likewise position priming was slightly stronger for old compared to young subjects but this effect too did not reach statistical significance (position priming strength: 13±1%

for 128 old subjects, 11±1% for 18 young subjects, $p = 0.18$, rank-sum test). As before position priming showed no clear difference between patients and age-matched controls (position priming strength: 14±1% for 10 patients, 13±1% for 57 age-matched controls, $p = 0.41$, rank-sum test).

**Do color and priming effects persist across trials?**

In the classic priming of pop-out studies, the authors reported a persistent effect of priming, whereby responses were faster even for trials that were preceded by the same color or position a few trials previously (Maljkovic and Nakayama, 1994, 1996). We therefore wondered whether the time course of perceptual priming would differ across subject groups. To this end, we calculated the priming strength for trials with same or different target color or position separated by increasing numbers of trials, without putting any conditions on the intervening trials. This revealed persistent priming effects for both color and position across many trials, with a time course that was longer for position (Figure S1B) compared to color (Figure S1A). However, there were no clear differences between groups except the ones described above: Color and position priming were significantly different in old compared to young subjects only for the immediately preceding trial (Figure S1A,C). We observed no consistent differences in the time course of color priming (Figure S1B) or position priming (Figure S1D) between patients and age-matched controls.

**Motor priming effects**

We wondered whether motor priming was different across the four groups. Motor priming did not differ between young and old subjects (priming strength: -1±1% for young, 0±0% for old, $p = 0.09$, rank-sum test) or between patients and age-matched controls (priming strength: 0±1% for patients, 1±0% for age-matched controls, $p = 0.55$, rank-sum test). But its incidence was different across age (percent of subjects with significant motor priming: 14% in young, 7% in old, $p < 0.05$, chi-squared test) and across cognitive impairment (0% in patients, 10% in age-matched controls, $p < 0.000005$, chi-squared test). Thus, motor priming is weakly present in general but declines across age and cognitive impairments.

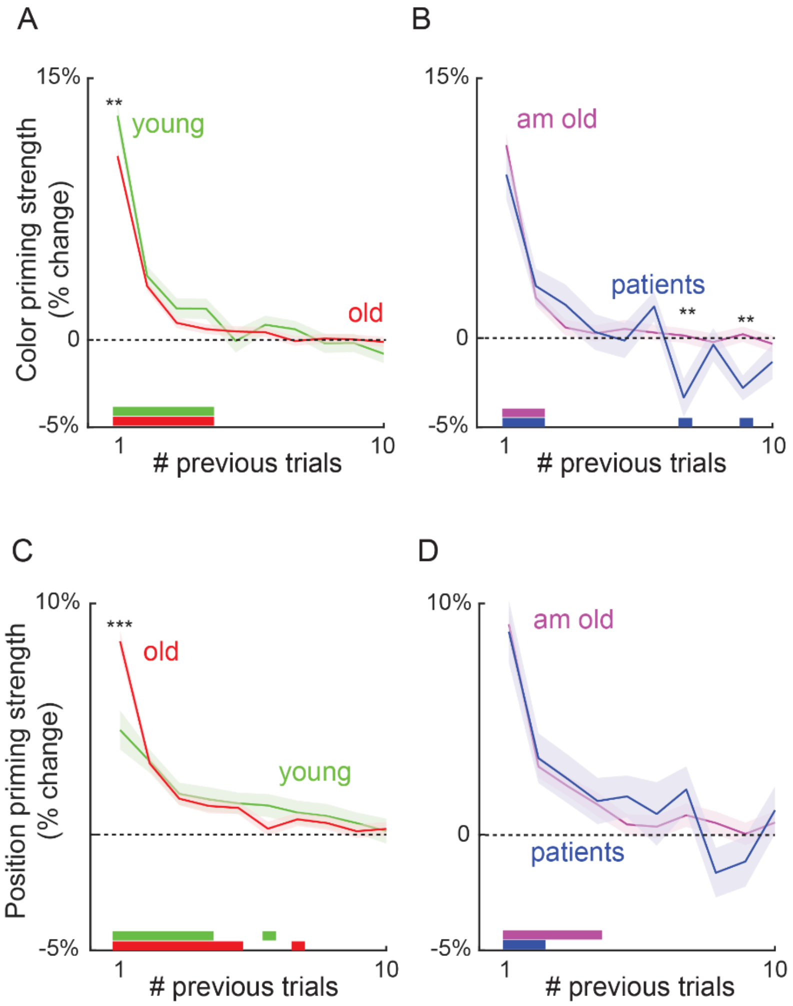


**Figure S1. Time course of perceptual priming effects**

(A) Time course of color priming strength as a function of number of previous trials, for old and young subjects. Asterisks indicate statistically significant difference between the priming strengths of the corresponding groups, using a rank-sum test across subjects. Shaded error bars indicate the s.e.m across subjects. Horizontal color bars of each color indicate time points at which the average priming strength of that group was significantly different from zero ($p < 0.05$ on a t-test on priming strength across subjects). Individual group priming strength significantly different from zero is indicated by color patches along the horizontal axis (green for young, red for old).

(B) Same as in (A) but for age-matched controls (purple) and patients (blue).

(C-D) Same as A & B but for position priming effects.
